# Neprilysin 1 in the Mushroom Body controls Feeding behavior by regulating SIFamide levels in *Drosophila melanogaster*

**DOI:** 10.64898/2026.10.10.758222

**Authors:** El-Sayed Baz, Daan Moons, Maria Pons Vizcarra, Veerle Vulsteke, Sofie De Groef, Isabelle Rahioui, Jiajun Zhang, Pedro Da Silva, Thomas Riemensperger, Federica Calevro, Patrick Callaerts

## Abstract

Feeding-related behavior in *Drosophila melanogaster* encompasses olfactory and gustatory detection of odors, foraging, and food intake. In *Drosophila melanogaster*, the neuropeptide SIFamide (SIFa), has been shown to integrate hunger signals to promote feeding in *Drosophila* and is considered a central regulator of mutually exclusive behavioral states. Here, we identify the mushroom bodies as a key downstream circuit of SIFamidergic neurons for regulation of food-seeking and feeding. Furthermore, we show that mushroom bodies are not merely downstream effectors but actively shape SIFamidergic signaling through the peptidase Neprilysin 1 for which SIFamide is a substrate. Thus, SIFa signaling is embedded within a feedforward-feedback circuit that couples internal state to the selection and execution of feeding behavior. We propose that the distinct neuronal subtypes postsynaptic to the elaborately arborized SIFamidergic neurons define discrete neuronal subroutines that differentially control the mutually exclusive behaviors related to sexual reproduction, sleep and feeding. Our findings reveal an evolutionary ancient peptidergic system that is modified by recruitment of novel genes for the flexible control of distinct behavioral programs and feeding strategies.

## INTRODUCTION

Feeding is crucial for survival in all animals, as it ensures energy homeostasis under varying nutritional conditions (Herzog, 2020; Mahishi et al., 2024; Münch et al., 2022). Feeding encompasses olfactory and gustatory detection of odors, food seeking or foraging, and food intake (Boesveldt & de Graaf, 2017; Mahishi et al., 2024; Martelli et al., 2017; McCrickerd & Forde, 2016; Sachse & Beshel, 2016; Stark, 2024; Zjacic & Scholz, 2022). This complex behavior is regulated by neuronal and endocrine systems that integrate metabolic needs with external stimuli (Hao et al., 2020; Mahishi et al., 2024). In both mammals and insects, neuropeptides play vital roles as orexigenic and anorexigenic signals that modulate feeding-related behaviors (Lin et al., 2019; Nässel et al., 2019; Root et al., 2011). In mammals, these signaling peptides are released by peripheral organs such as the adipose tissue, pancreas, and gastrointestinal tract, and their effects are primarily integrated by the hypothalamus (Chen et al., 2025; Sobrino Crespo et al., 2014). In insects, a similar system conveys information about metabolic status through neuropeptides that is subsequently integrated in the brain (Audsley & Weaver, 2009; Nagata & Zhou, 2019; Nässel, 2025; Nässel et al., 2019; Nässel & Zandawala, 2020). In *Drosophila melanogaster*, several neuropeptides have been implicated in encoding hunger and satiety states (Bhumika & Singh, 2018; Lin et al., 2019; Pool & Scott, 2014; Wang & Wang, 2019). For example, neurons secreting Neuropeptide F (NPF) mediate a feeding response by integration of both food attractiveness and hunger state (Fadda et al., 2019; Lee et al., 2004; Shen & Cai, 2001). Short NPF modulates olfactory sensitivity to nutrient-related odors, thereby influencing feeding (Fadda et al., 2019; Lee et al., 2004, 2009). Hugin-producing neurons are responsible for suppressing immediate feeding responses during satiety (Itskov & Ribeiro, 2013). Insulin-like peptides and adipokinetic hormone are involved in regulating feeding and foraging behaviors (Jourjine et al., 2016; Liessem et al., 2023; Sudhakar et al., 2020). Drosulfakinin, allatostatin A, myoinhibitory peptide and SIFamide are additional *Drosophila melanogaster* neuropeptides that have been implicated in feeding (Chen et al., 2016; Hergarden et al., 2012; Kubrak et al., 2022; Min et al., 2016; Söderberg et al., 2012).

SIFamide (SIFa) has been shown to translate hunger signals into feeding behavior. Specifically, activation of SIFamidergic neurons increases appetitive olfactory and gustatory behavior as well as food intake (Martelli et al., 2017). The SIFamide receptor (SIFaR) (Jørgensen et al., 2006) and the vertebrate gonadotropin inhibitory hormone receptor (GnIHR) are homologs (Ubuka & Tsutsui, 2014), although their respective ligands, SIFa and GnIH, are not sequence-related (Jékely, 2013; Ubuka & Tsutsui, 2014). The GnIHR integrates metabolic information to regulate the allocation of behavioral priorities between food intake and reproduction, favoring feeding during metabolic stress (Clarke et al., 2012; Smith & Clarke, 2010). GnIH, the ligand of the GnIHR, is produced in the hypothalamus (Lee et al., 2008; McConn et al., 2014). SIFa, the ligand of the SIFaR, is mainly produced by four neurons in the pars intercerebralis of all insects studied to date (Arendt et al., 2016; Janssen et al., 1996; Ons et al., 2011; Terhzaz et al., 2007; Verleyen et al., 2004, 2009). The pars intercerebralis of insects has been described as the functional equivalent of the hypothalamus (de Velasco et al., 2007; Hartenstein, 2006; Tessmar-Raible, 2007). These parallels have led to the hypothesis that GnIH and SIFa, along with their receptors, have evolved from a common ancestral signaling system (Jékely, 2013; Mirabeau & Joly, 2013; Ubuka & Tsutsui, 2014).

SIFa is highly conserved across arthropods (Verleyen et al., 2004). It was first isolated from the grey flesh fly *Neobellieria bullata* where it was named leucopyrokinin (Janssen et al., 1996) and subsequently identified in numerous insect species (Gellerer et al., 2015; Hauser et al., 2006; Jørgensen et al., 2006; Junhong et al., 2023; Rahman et al., 2013; Šimo et al., 2009; Verleyen et al., 2004, 2009). SIFa has been shown to modulate several behaviors, namely sleep, sexual behavior, and feeding. In *D. melanogaster*, SIFa plays an important role in promoting sleep, although other, as yet unidentified neurotransmitters released by SIFamidergic neurons are also required for this effect. The SIFaR regulates sleep and feeding in function of the time of day (Huang et al., 2021; Park et al., 2014; Velazquez et al., 2025). SIFa also contributes to the control of sexual behavior. Ablation of SIFa neurons or knockdown of SIFaR in *fruitless*-expressing neurons results in indiscriminate courtship behavior towards both sexes (Sellami & Veenstra, 2015; Terhzaz et al., 2007). Moreover, SIFamidergic neurons control the balance between mating duration and feeding in males (Song et al., 2025). Finally, SIFa promotes appetitive behavior and hunger-dependent modulation of odor responses in *Drosophila* (Martelli et al., 2017), it exerts neuromodulatory control over the stomatogastric ganglion in the lobster *Homarus americanus* (Christie et al., 2006), has been proposed to regulate salivary gland secretion in the blood-feeding tick *Ixodes scapularis* (Šimo et al., 2009; Šimo et al., 2013), and promotes food intake in the Chagas disease vector *Rhodnius prolixus* (Ayub et al., 2020). Together, these studies are suggestive of anevolutionary conserved role for SIFa in the regulation of feeding-related behavior across arthropods.

SIFamidergic neurons integrate multiple signals that collectively reflect the internal state of the animal (Martelli et al., 2017; Zhang & Kim, 2026). SIFamidergic neurons in turn are elaborately arborized and form numerous postsynaptic contacts throughout the brain (Martelli et al., 2017). Martelli et al. (2017) identified functional connections in the antennal lobe that regulate olfactory signaling. In addition, anti-SIFamide immunoreactivity has been detected in the dorsal fan-shaped body, the optic lobes, mushroom bodies, suboesophageal ganglion and around the esophagus. The functional significance of these putative synaptic contacts in feeding behavior remains unknown. However, the fact that the mushroom bodies have – besides their role in learning, memory, and sensory processing – also been implicated in feeding behavior, identifies them as a compelling candidate for mediating at least some of the effects of SIFa in feeding behavior regulation (Akalal et al., 2006; Çoban et al., 2024; Heuer et al., 2012; Hige, 2018; Lin et al., 2014; McGuire et al., 2001; Tsao et al., 2018; Turner et al., 2008),

Once released by presynaptic neurons, neuropeptide signaling must eventually be terminated, in part through peptide degradation (Isaac et al., 2009). This is also true for the neural control of feeding given its dependence on neuropeptide-mediated neuromodulation (Audsley & Weaver, 2009; Chen et al., 2025; Hallier et al., 2016; Shioda et al., 2008; Taghert & Nitabach, 2012). Peptidases play key roles in the degradation of neuropeptides. Among these enzymes, neprilysins (NEPs) are known to terminate neuropeptide signaling at the cell surface in the brains of both mammals and insects (Hallier et al., 2016; Isaac et al., 2007; Isaac & Nässel, 2003; Sitnik et al., 2014; Siviter et al., 2000). Sitnik et al. (2014) have characterized Neprilysins 1-5, which had previously been identified as the *Drosophila* neprilysins most closely related to mammalian neprilysin (Meyer et al., 2021; Turner et al., 2001). Neprilysin 1 and 2 are both expressed in the pars intercerebralis of the adult brain and Neprilysin 1 is additionally expressed in the mushroom bodies (Sitnik et al., 2014), where it has been shown to be involved in short-term and long-term memory (Turrel et al., 2016).

In the current study, we identify a role for SIFamide in food-seeking behavior and in the microstructure of food intake. We next demonstrate that the SIFamidergic neurons form synaptic contacts with the mushroom bodies where SIFamide receptor is required for the control of these behaviors. Finally, we identify Neprilysin 1 as the peptidase responsible for switching off SIFamide signaling in the mushroom bodies by cleaving the peptide. These results demonstrate that SIFamide is involved in multiple, if not all, aspects of feeding behavior from food seeking to feeding bursts, that the mushroom bodies are postsynaptic targets of SIFamidergic neurons, and that Neprilysin 1 serves as an important molecular off-switch for SIFamidergic signaling.

## MATERIALS AND METHODS

### Fly strains and rearing

*Drosophila* strains were reared on standard cornmeal food (Table S1) in incubators (MLR-351, Sanyo, Japan) at 23 °C and 60% humidity under a 12:12-hour light/dark cycle. The fly stocks used in this work were Nep1^EY21255^ (BDSC #22465), UAS-rpr (BDSC #5824), SIFaR (RNAi) (BDSC #34947), SIFaR (RNAi) (BDSC #44068), CRIMIC-TG4 Nep1 (BDSC #79257), Nep1 RNAi (BDSC #64911); Nep1 RNAi (BDSC #65960), Nep1 RNAi (VDRC #27537), Nep1 RNAi (VDRC #108660), OK107-GAL4 (gift from Sean Sweeney), SIFaR-Gal4 (BDSC #76670), SIFamide-Gal4 (gift from Young-Joon Kim), MB247-Gal4 (gift from Sha Liu), UAS-CD8GFP (BDSC #5137), tub-Gal80^ts^ (BDSC #7019), Nep1^Ey21255^ revertant (this study); UAS-Nep1 (this study), Wildtype Canton-S (Ron Davis), wildtype iso31 (BDSC #5905) orco^2^ (BDSC #23130). Parental crosses were set up in *Drosophila* culturing bottles containing 50 mL of food. After 3 days, the parents were removed, and the progeny allowed to develop to adulthood.

### Production of Nep1 overexpression flies

A clone (GH03315) containing full-length Nep1 cDNA was obtained from the Drosophila Genomics Resource Center (Indiana University). The clone was digested with *Ndel* and the purified Nep1 fragment was ligated in the *Xbal* restriction sites of the pUAST vector (Brand & Perrimon, 1993). The resulting construct was sequence verified. Transgenic flies were generated as a service by Duke University Model System Genomics.

### Production of the Nep1^EY21255^ revertant line

A precise excision of the Nep1^EY21255^ P-element insertion following the protocol described by Hummel & Klämbt (2008). Progeny that lost the *white^+^* marker were used to establish independent stocks and the nature of the excision was determined by means of PCR and sequencing.

### Behavioral assays

#### Food-seeking behavior

The hunger-driven food-seeking behavior for single fly assay was based on (Tsao et al., 2018). Newly emerged adult flies were transferred to food vials and kept for 3 days. Groups of 10-15 mated female flies were selected under CO_2_ anesthesia and allowed to recover for about 2 days on fresh standard fly food. Twenty-four hours before the experiments, the flies were transferred without anesthesia into vials containing 1% agar (LP0013, Oxoid, England) instead of regular food to induce the hunger state. During the testing phase, each 5-7-day-old fly was transferred from the starvation vial and introduced near the wall of a Petri dish (628102, Greiner Bio-One, Frickenhausen, Germany, 60 mm diameter and 6 mm height) using a mouth aspirator. Individual flies were allowed to move freely in the Petri dish for 10 minutes to locate a small drop (5 µl) of yeast food placed in the center of each dish. The yeast food solution was prepared by mixing 2% (w/v) dry yeast [200 mg, 5406105, Algist Bruggeman, Gent, Belgium] and 10% (w/v) sucrose (1 g, 50389, sigma-Aldrich) in 10 ml of Baxter water [AKF7114, Baxter, Zurich, Switzerland], followed by 16 h of incubation at 28 °C in a shaking incubator (170 rpm) [C25 New Brunswick scientific, Edison, NJ, USA]. A fly was considered to have located the food drop after remaining on it for at least 3 seconds. This criterion was used to exclude flies that accidentally passed over or briefly contacted the yeast drop (Tsao et al., 2018). The hunger-driven food-seeking performance was calculated based on how quickly starved flies found the yeast drop in a 10-minute session. The food-seeking index was calculated as follows: [Total assay time (600 sec) – the time taken to find food (sec)]/Total assay time (600 sec). The index ranged from 0 to 1, with values closer to 1 indicating rapid food localization, and values closer to 0 indicating impaired food-finding. If a fly explored the Petri dish but failed to locate the food drop within 600 seconds, a food-seeking index of 0 was assigned, and the trial was terminated. Per session, 15 new Petri dishes were monitored simultaneously. Experimental groups and their corresponding controls were randomly distributed across dishes within each session and experiments were repeated across different days using progeny from at least four independent parental crosses. All sessions were performed between ZT1 and ZT3 in a temperature and humidity-controlled behavioral cubicle/chamber (∼23° C, 60-80% relative humidity) equipped with dimmable red LED light strips (SMD3528-IP67, LEDLightsWorld), to allow testing under dim red light to minimize visual inputs to the fly. Flies were placed in the behavioral room for 1 hour prior to testing.

#### FlyPAD

Feeding behavior was assessed using the fly Proboscis and Activity Detector (flyPAD) system (Easy Behavior, Lisbon, Portugal), which enables high-resolution quantification of feeding events through capacitive sensing (Itskov et al., 2014). The system consists of electrodes that detect changes in capacitance when a fly contacts the food source via its proboscis allowing individual feeding events to be recorded. For each experiment, 5 µL of food was loaded into one well of the flyPAD arena. The food consisted of 2% yeast, 1% agar, and 10% sucrose. The second well was left empty but sealed from below with tape to prevent escape while maintaining the structural integrity of the arena. All experiments were conducted between ZT1 and ZT3 to control for circadian influences on feeding behavior. Flies were loaded into the flyPAD arenas in a completely dark room under dim red light illumination. Dim red light conditions were used throughout handling and experimentation to minimize visual cues and ensure that behavior was primarily guided by non-visual sensory modalities, such as olfaction. The flyPAD system was operated inside an incubator maintained at 22 °C, also under constant redlight conditions, to ensure stable environmental parameters throughout the experiment. Prior to each experimental session, the recording system (Bonsai software) was checked to ensure that all sensors were functioning optimally. Flies were introduced individually into the arenas using an aspiration device and allowed to adapt for 15 minutes before the start of recording. Feeding behavior was subsequently recorded for 1 hour. Capacitive signals corresponding to feeding events were acquired using the flyPAD system and processed via Bonsai software for downstream analysis as per the manufacturer’s instructions.

### Antibody production

Full-length recombinant His-Neprilysin1 was produced by the VIB Protein Service Facility (PSF) as a service. Purified recombinant protein was used to generate polyclonal antibodies in rabbits using the speedy 28-day program of Eurogentec. The IgG fraction was isolated from the serum with the Thiophilic Adsorbent (Pierce, Life Technologies) following the manufacturer’s instructions.

### Immunohistochemistry

Following dissection in icecold 1x PBS, adult fly brains were fixed in 2% formaldehyde for 50 min. Brains were then washed three times in PBS containing 0.5% Triton X-100 (PBST) prior blocking in PAXD (PBS containing 1% BSA, 0.3% Triton X-100, 0.3% deoxycholate) for 30-60 minutes. Subsequently, brains were incubated overnight at 4°C with primary antibodies in PAXD, washed at least three times for 15 minutes each with 0.5% PBST, and incubated overnight at 4°C with secondary antibodies in PAXD. Brains were then washed in 0.5% PBST, mounted in Vectashield mounting medium (Vector Laboratories Cat #H-1000), and imaged with an Olympus FluoView FV1000 confocal microscope. Image stacks were processed in Fiji ImageJ (NIH) and Photoshop software (Adobe). Primary antibodies used in this study include mouse α-Bruchpilot (Brp) (DSHB, nc82, 1:30), mouse α-GFP (Roche 11814460001, 1:1000), rabbit α-GFP (Life Technologies, Ref#: A6455; 1:1000), rabbit anti-SIFamide (gift from Jan Veenstra, Bordeaux, 1:2500), and rabbit anti-Nep1 (this study, 1:100). Secondary antibodies were donkey α-mouse Alexa-488, donkey α-mouse Alexa-594, and donkey α-rabbit Alexa-594. All secondary antibodies were from Jackson ImmunoResearch and were used at a 1:200 concentration.

### Neprilysin-1 activity assay

For the neuropeptide enzymatic assay, we generated a stable S2-cell line expressing Nep1 under the control of an inducible promoter. Nep1 expression and the optimal induction time were determined using western blot analysis. This analysis also confirmed the absence of endogenous Nep1 expression in control S2 cells. Neprilysin activity was assessed using the selective fluorescent substrate Mca-RPPGFSAFK (Enzo Life Sciences). Experiments were performed at 25°C in 24-well plates using approximately 6 x 10^6^ cells per well in a final volume of 100 μm of medium. The substrate was added at a final concentration of 20 μM to control (S2 cells) and Nep1 cells (Nep1 stable cell line) at different time points (30, 60, and 90 min). Phosphoramidon (Enzo Life Sciences) and Thiorphan (Enzo Life Sciences) were used as specific inhibitors for Neprilysin. Samples with protease inhibitors (10 μM) were pre-incubated for 30 min before adding the substrate. The blank sample consisted of cells incubated in medium without treatment. The amount of cleaved substrate was determined by comparing the fluorescent signal at 400 nm following excitation at 320nm. These experiments confirmed that Nep1 is inducibly expressed in S2 cells and exhibits enzymatic activity. This assay was subsequently used to determine whether Nep1 can cleave SIFa neuropeptide, thereby identifying the latter as a substrate. For this experiment, approximately 6 × 10⁶ cells were used per well in a final volume of 100 μL of medium at 25°C. The inhibitor phosphoramidon (Enzo Life Sciences) was added 30 min before adding the SIFa neuropeptide at a final concentration of 10μM. The supernatant was collected at 4 different time points (0, 15 min, 30 min, and 60 min) for a total of three replicates at each time point with and without the inhibitor. The only time point not tested with inhibitor is time 0min. The blank sample consisted of cells with medium without treatment.

### HPLC assays

Cleavage of physiological peptides was assessed by monitoring the disappearance of the parent peptide by reverse-phase high-performance liquid chromatography (HPLC), following the principle of a protocol previously used to assess the cleavage specificity of neprilysin (Sexton et al., 2012). Reactions were run in triplicate. HPLC was carried out using a Synergi C12 4 µm Max-RP 80 Å, LC Column (250 x 4.6) using a linear gradient from 0.1% trifluoroacetic acid (TFA) in 95% water and 5% acetonitrile to 0.1% TFA in 50% acetonitrile/water. Peptides were eluted at 1 ml/min with a 20 min gradient from 5% to 50% acetonitrile in water (0.04% TFA), detected at 214 nm and quantified by measuring the corresponding peak areas.

### Connectome data analysis

To investigate the interaction between SIFamidergic neurons and mushroom body (MB) circuits, we analyzed hemibrain connectome data from the *Drosophila melanogaster* adult female brain using the neuPrint (https://neuprint.janelia.org) and FlyWire (https://flywire.ai) platforms. Neuron types were identified and selected based on established morphological and neurotransmitter identity criteria, using both neuPrint annotations and cross-referenced published classifications. To construct the connectivity diagram, we extracted all postsynaptic partners of the SIFamidergic neurons within the MB region, with particular attention to neurons forming direct or indirect connections within MB-associated circuits. These included anterior paired lateral (APL) neurons, dorsal paired medial (DPM) neurons, PAM04 dopaminergic neurons targeting the β2 lobe, lateral horn output neurons (LHONs), and identified mushroom body output neurons (MBONs) from specific compartments (γ1,2; γ4, γ5; α2). Synaptic sites were determined using the 5-synapse threshold suggested by FlyWire to exclude false positives (Dorkenwald et al., 2024).

### RNA extraction, cDNA synthesis, and qRT-PCR

Total RNA was extracted using the NucleoZOL one-phase RNA purification kit (Macherey-Nagel, Düren, Germany). Briefly, fly heads were homogenized in 500 µL of NucleoZOL using the FastPrep-24™ 5G bead beating system (MP Biomedicals, USA). After homogenization, 200 µL of RNase-free water was added, and the mixture was vigorously shaken by hand. Samples were incubated at room temperature for 5 minutes and then centrifuged at 12,000 rcf for 15 minutes. From the resulting supernatant, 500 µL was transferred to a fresh RNase-free 1.5 mL tube containing 500 µL of 99.5% isopropanol. The samples were gently inverted to mix, incubated for 10 minutes at room temperature, and centrifuged at 12,000 rcf for 10 minutes at 4 °C. The RNA pellet was washed twice with 500 µL of 75% ethanol and resuspended in 16 µL of RNase-free water. RNA concentrations were measured using a NanoDrop ND-1000 spectrophotometer (Thermo Fisher Scientific, Wilmington, USA), and samples were stored at −80 °C until further use.

First-strand cDNA was produced using the SensiFAST™ cDNA Synthesis Kit (GC Biotech, Waddinxveen, The Netherlands) in a 20 µL reaction containing 1 µg of total RNA, according to the manufacturer’s protocol. The resulting cDNA was diluted in nuclease-free water to a final concentration of 100 ng/µL and stored at −20 °C. Quantitative real-time PCR (qRT-PCR) was performed using the SYBR Green system (FastGene 2× IC Green mix—low ROX, Nippon Genetics, Düren, Germany). Each 10 µL reaction contained 5 µL SYBR Green mix, 1 µL primer mix (5 µM each forward and reverse), 2 µL nuclease-free water, and 2 µL cDNA. Reactions were performed in duplicate using 384-well optical plates using a QuantStudio 6 Pro Real-Time PCR System (Thermo Fisher Scientific, USA). Relative transcript levels of Nep1 were quantified using the ΔΔCt method and normalized to two reference genes, LaminCa and Actin42A (De Groef et al., 2024). Gene-specific primers were designed using NCBI Primer-BLAST.

Primer sequences were:

Nep1 Fw: ATG AAC GCG ACA ACA AAG CC

Nep1 Rv: ATT TGA GGA ATA TCC ATG CAG CTC

LamCa Fw: TCC ACC CAA CAA TCT GGT GAT

LamCa Rv: GCA ACA TCC TCT TTG TCG GC

Act42a Fw: TGC AAA AGG AAA TCA CGG CG

Act42a Rv: CCG CCG ATC CAA ACA GAG TA

Expression data were analyzed and visualized using GraphPad Prism version 10.4.1 (GraphPad Software Inc., San Diego, USA).

### Statistical analysis

Data were analyzed using Prism software version 10.4.1 (GraphPad Software Inc., San Diego, USA). The data were assessed for normality using the Kolmogorov-Smirnov test. One-way analysis of variance (ANOVA) with Tukey’s post hoc test for normally distributed data, Kruskal–Wallis ANOVA test with Dunn’s post hoc test for nonparametric data. Comparison between two groups were tested with Mann Whittney test. For HPLC data, unpaired or paired t-tests were performed to compare the two treatments or within the same treatment, respectively. Error bars represent the standard error of the mean (SEM).

## RESULTS

### SIFamide regulates hunger-driven food-seeking and feeding microarchitecture

SIFa has previously been shown to translate hunger signals into feeding behavior in *Drosophila* (Martelli et al., 2017). Specifically, activation of the SIFamidergic neurons causes satiated flies to exhibit behaviors normally observed in starved flies, including increased attraction to the food-associated odor ethylacetate, an increased proboscis extension reflex, and increased food intake in the CAFE assay (Martelli et al., 2017). Because yeast is a major nutritional source for *Drosophila* species (Scheidler et al., 2015), we sought to examine the role of SIFa neurons in hunger-driven innate food-seeking behavior (foraging) using yeast as an attractant. In addition, since hunger has been shown to affect the microstructure of feeding (Itskov et al., 2014), we hypothesized that SIFa might contribute to this hunger-dependent modulation.

To maximize the potential effects of SIFa loss, we ablated SIFa neurons using UAS-rpr (*Reaper*) and the SIFa-GAL4 driver line. We first validated the SIFa-GAL4 line by comparing its expression pattern with SIFa peptide localization (Figure 1A, B). The results showed four SIFamidergic neurons, organized in two pairs with large somata situated in in the pars intercerebralis, consistent with previously published observations (Tehrzaz et al., 2007) and validate SIFa-GAL4 as completely reproducing endogenous SIFa expression.

**Figure 1.**
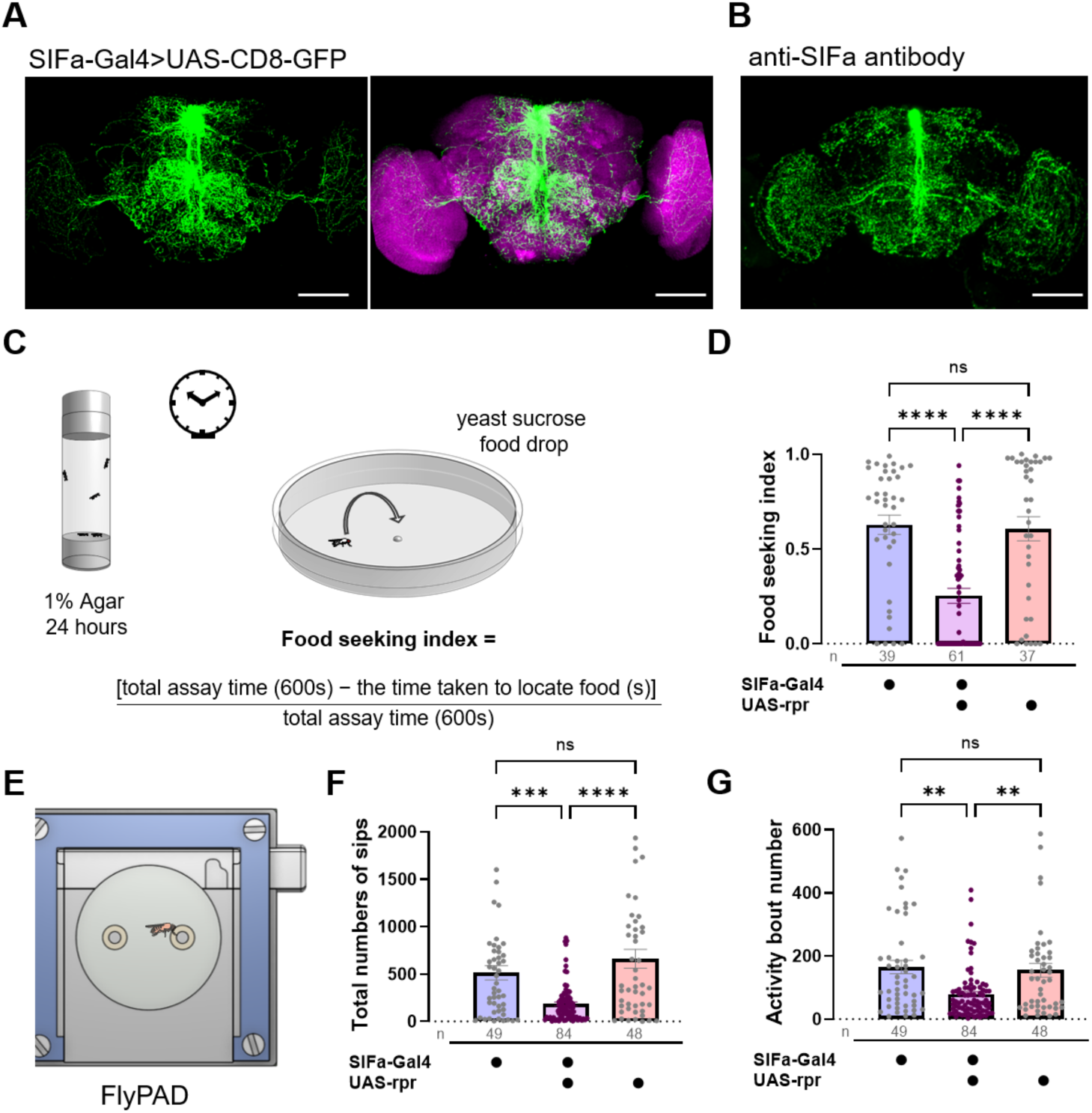
Extended food seeking and altered feeding microstructure in flies with ablation of SIFamide neurons. **(A)** SIFa expression pattern on adult SIFa-Gal4>UAS-CD8-GFP brain. SIFa is produced in only two pairs of neurons with large stomata that are situated in the pars intercerebralis but have extensive arborization in the brain. The size bar is 100 µm. **(B)** WT flies were stained against SIFamide (anti-SIFa, rabbit antibody) to determine its expression pattern The size bar is 100 µm. **(C)** Schematic representation of the behavioral assay of hunger-driven food-seeking behavior. Flies were deprived of food for 24 hours and placed in vials containing 1% to induce a hunger state. During the testing phase under dim red light, single-starved flies were transferred and allowed to move freely within each Petri dish for 10 minutes to locate a small drop (5 µl) of yeast food placed in the center of each dish. The food-seeking performance was calculated based on how quickly starved flies found the yeast drop in a 10-minute session. Indexes were ranged between zero and one, with values closer to one indicating that the food was located quickly, while values closer to zero reflect impaired food-finding performance. **(D)** SIFamide>rpr flies display impaired food-seeking behavior in comparison to parental controls. Kruskal-Wallis tests followed by Dunn’s multiple comparisons test. **(E)** Schematic representation of the behavioral assay of fly Proboscis and Activity Detector (flyPAD) system. **(F)** SIFamide>rpr flies display fewer numbers of sips in comparison to parental controls. Kruskal-Wallis tests followed by Dunn’s multiple comparisons test. **(G)** The Activity bout number was lower in flies with ablation of SIFamide neurons. Kruskal-Wallis tests followed by Dunn’s multiple comparisons test. Plots represent mean ± SE, and each dot represents data measurement for one individual. Number of tested flies represented below each bar for each genotype. Asterisks represent statistical significance, \*\**p* < 0.01, \*\*\**p* < 0.001, \*\*\*\**p* < 0.0001, ns, not significant.

Ablation of SIFa neurons in the brain resulted in a significantly reduced food seeking index compared with control flies (Kruskal-Wallis: H = 34.71, df = 2, *p* < 0.0001) (Figure 1C, D). The reduced food-seeking index reflects a longer latency to locate the yeast drop in SIFa neuron-ablated flies, with a subset of flies failing to locate the yeast within the time allotted for the experiment.

Furthermore, flies lacking SIFa neurons exhibited significantly fewer sips and activity bout number than control flies (sips: Kruskal-Wallis: H = 20.84, df = 2, p < 0.0001; activity bout: Kruskal-Wallis: H = 16.35, df = 2, p < 0.0003) (Figures 1F). Because sip number has been shown to correlate linearly with food intake (Itskov et al., 2014), this reduction is consistent with a decrease in food intake.

Together, these results demonstrate that SIFamidergic neurons are required for hunger-driven innate food-seeking behavior when yeast serves as the food-associated cue. In addition, they show that SIFamidergic neurons influence feeding behavior by regulating its microstructure.

### The SIFa receptor is required in mushroom bodies to control food seeking behavior and regulate feeding microstructure

To date, a role for the SIFa receptor (SIFaR) in reproductive behavior and sleep has been demonstrated using RNAi-mediated knockdown (Cavanaugh et al., 2014; Park et al., 2014; Sellami & Veenstra, 2015). Complete loss of the SIFaR confirmed the role in sleep and impacted locomotion and feeding in a time-of-day specific manner (Velazquez et al., 2025). The SIFaR has been reported to be expressed in many neurons (Dreyer et al., 2019; Velazquez et al., 2025), raising the question of which neuronal populations require the receptor to control food-seeking behavior and regulate feeding microstructure. Given that the mushroom bodies have been shown to regulate food-seeking behavior by integrating hunger and satiety signals (Tsao et al., 2018), we hypothesized that expression of SIFaR in the mushroom bodies would be required in food seeking and feeding microstructure. To test this hypothesis, we first examined SIFaR expression in the mushroom bodies using a SIFaR GAL4 insertion line (Figure 2A). GFP expression driven by SIFaR-GAL4 was widespread throughout the brain with prominent expression in the mushroom bodies. We next analyzed the connectome of SIFamidergic neurons. Two of these neurons were found to project to the MB region. For each hemisphere, we quantified synaptic contacts specifically onto Kenyon cells (KCs), the intrinsic neurons of the MB, and identified 34 input synapses on the left hemisphere and 23 on the right (Figure 2B). In addition, SIFamidergic neurons also formed synaptic contacts with other MB circuit neurons (see Suppl. Figure S1). SIFa signaling may influence mushroom body function through both synaptic transmission and volume transmission. Given the prominent expression of SIFaR in the mushroom bodies and the direct synaptic input from SIFamidergic neurons onto KCs, we focused on the requirement for SIFaR in mushroom body intrinsic neurons for the regulation of feeding behavior. Targeted RNAi-mediated knockdown of the SIFa receptor in these neurons using the specific GAL4 driver line MB247 significantly reduced the food-seeking index. The same phenotype was observed with two SIFaR-RNAi effector lines: line 34947 (Kruskal-Wallis: H = 14.19, df = 2, *p* = 0.0008), and line 44068 (Kruskal-Wallis: H = 11.02, df = 2, *p* = 0.004) (Figure 2C). In addition, significant reductions in number of sips and feeding activity bouts were observed compared to the parental controls (Figure 2D, 2E). These results demonstrate that SIFaR in the mushroom body intrinsic neurons is required for normal innate food-seeking behavior and contributes to the regulation of feeding microstructure.

**Figure 2:**
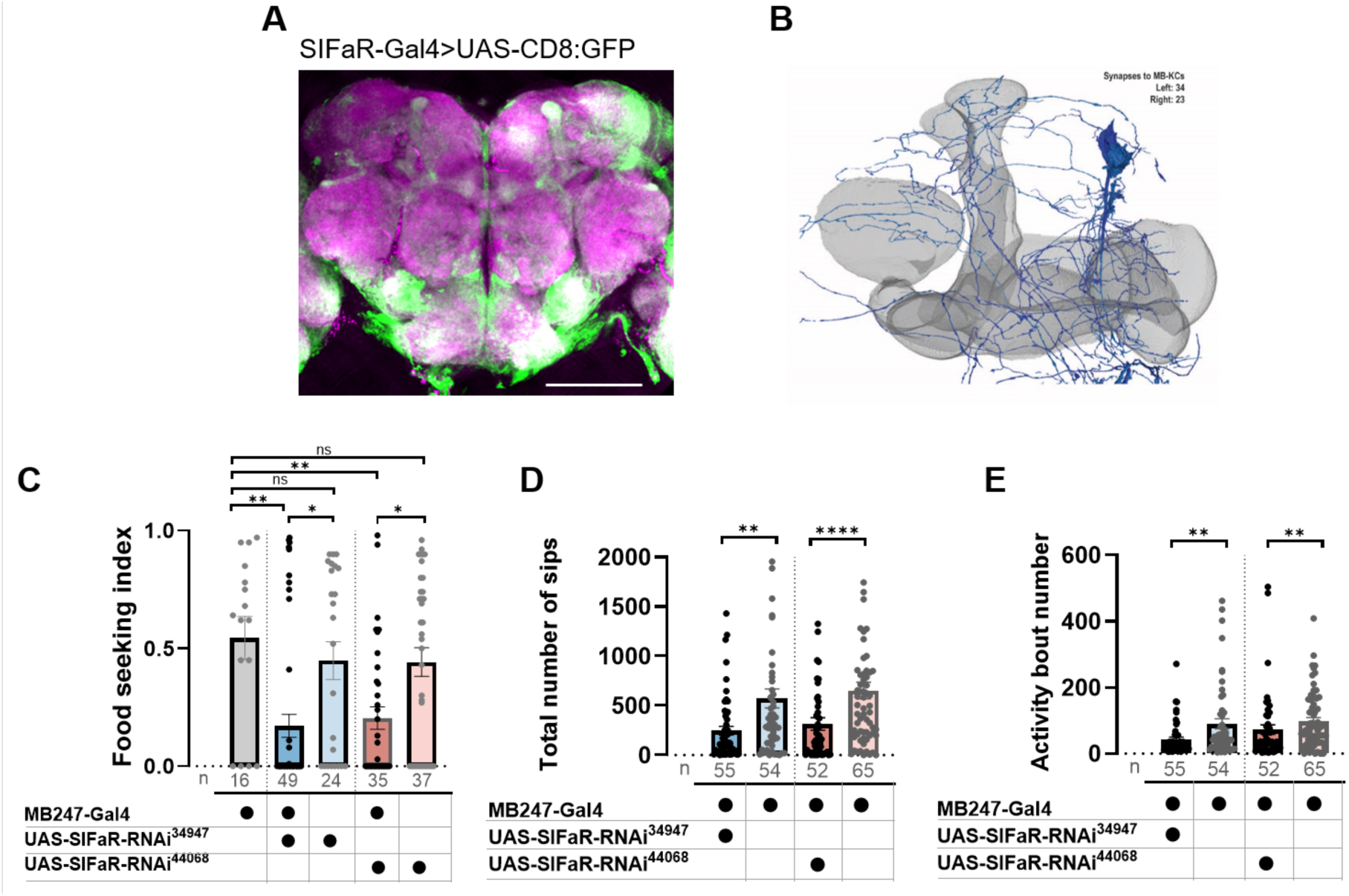
SIFa signaling in the mushroom body controls hunger-driven food-seeking behavior and food intake. **(A)** SIFa receptors (SIFaR) are expressed broadly throughout the adult brain, with notably high levels in the AL and MB (SIFaR-Gal4>UAS-CD8:GFP). The size bar is 100 µm. **(B)** Interaction of SIFamidergic neurons and mushroom body circuits: Two of the four SIFa neurons (blue) send projections to the mushroom body (grey) of each brain but only form very limited connections to Kenyon cells (34 input synapses on the left and 23 on the right brain side). **(C)** Knockdown of SIFa receptor in the MB impaired hunger-driven food-seeking behavior using SIFaR-RNAi 34947 and 44068 effector lines. Kruskal-Wallis tests followed by Dunn’s multiple comparisons test. **(D)** The total number of sips was lower when the SIFa receptor was knocked down in the MB using SIFaR-RNAi effector lines 34947 and 44068 compared to parental controls. Mann-Whitney U test. **(E)** The Activity bout number was lower when the SIFa receptor was knocked down in the MB. Mann-Whitney U test. Plots represent mean ± SE, and each dot represents a data measurement for one individual. The number of tested flies is represented below each bar for each genotype. Asterisks represent statistical significance: *p < 0.05, **p < 0.01, ***p < 0.001, ****p < 0.0001, ns, not significant.

### Neprilysin 1 is expressed in the mushroom bodies and degrades SIFa

After showing that SIFamidergic neurons control innate food-seeking behavior and feeding microstructure via SIFaR in the mushroom intrinsic neurons, we explored how the signal is terminated. SIFa signaling needs to be inactivated either by receptor desensitization and internationalization or through degradation of the peptide ligand. Given the demonstrated role of neprilysins in inactivation of peptide signaling, we hypothesized that a neprilysin would be responsible. We focused on Neprilysin 1, the only *D. melanogaster* neprilysin that has previously been shown to be expressed in the adult mushroom bodies by means of in situ hybridization (Sitnik et al., 2014).

We first analyzed the distribution of Nep1 in the adult brain using a CRIMIC-GAL4 insertion in the Nep1 locus to drive GFP expression and by a newly generated polyclonal anti-Nep1 antibody. The GFP reporter assay revealed prominent Nep1 expression in the mushroom bodies and antennal lobes, with additional expression in the optic lobes and the pars intercerebralis (Figure 3A). Immunohistochemistry with anti-Nep1 antibody showed strong staining in the mushroom bodies and antennal lobes (Figure 3B), confirming that Nep1 is strongly expressed in the mushroom bodies and antennal lobes.

**Figure 3.**
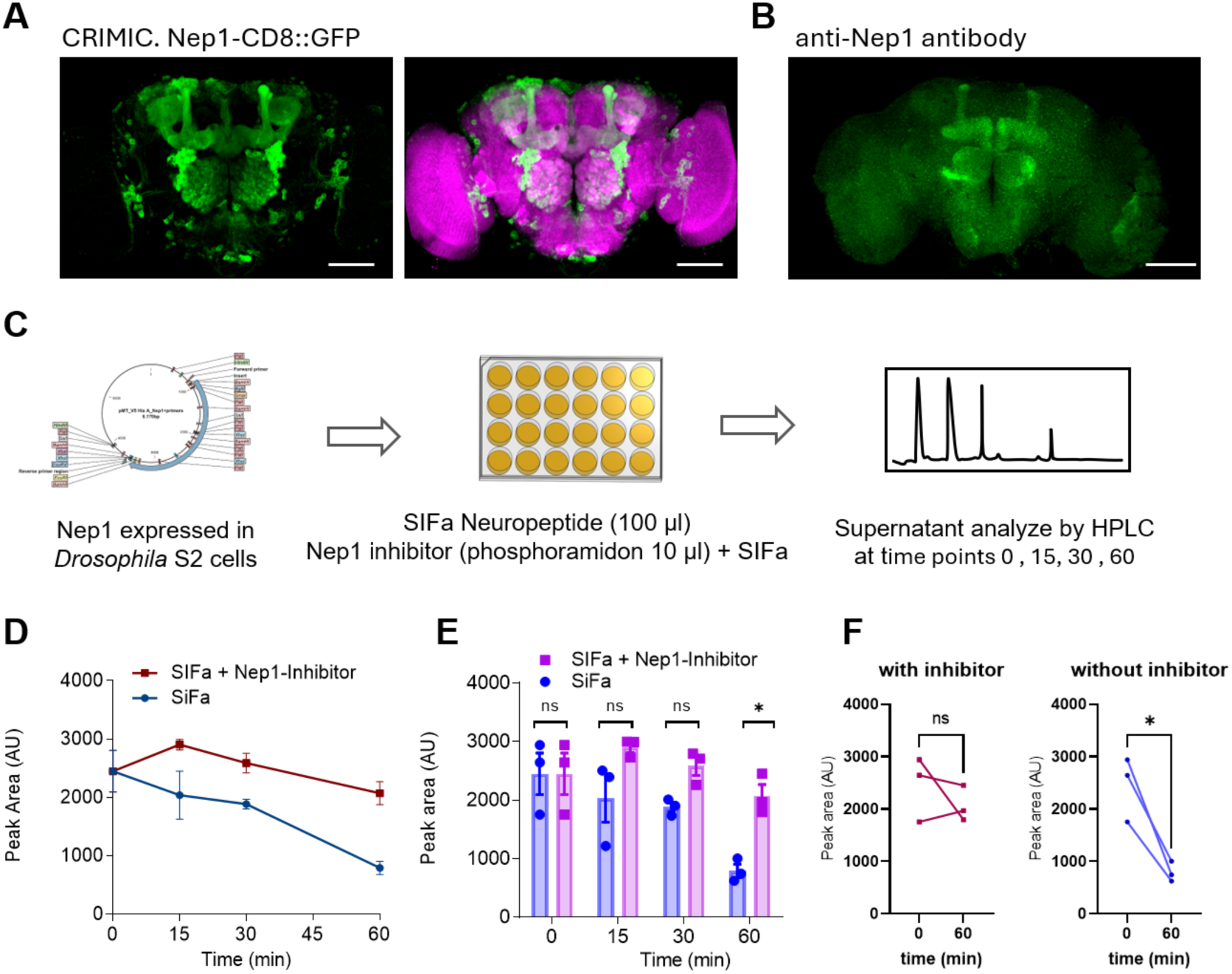
SIFamide is a substrate of Nep1. (**A-B**) Neprilysin 1 expression pattern in *Drosophila* adult brain. **(A)** Nep1 expression pattern driven by CRIMIC CD8::GFP. Nep1 is enriched in the mushroom bodies and antennal lobes, with additional expression in the cell bodies of the optic lobes and pars intercerebralis of the *Drosophila* adult brain. **(B)** WT flies were stained against Nep1 (anti-Nep1 (IgG mix) rabbit antibody) to determine its expression pattern. Cy3 was used as the secondary antibody. (A) General view of the Drosophila brain where Nep1 expression is seen in the mushroom bodies and the antennal lobes. The size bar is 100 µm. **(C)** Schematic illustration for Neprilysin-SIFamide activity assay. A stable S2-cell line expressing Nep1 was used to test whether Nep1 can cleave the SIFa neuropeptide, determining whether it is a substrate. Experiments were performed in a 24-well plate. Around 6 million S2 cells and the Nep1 stable cell line were used for each assay. The Nep1 inhibitor phosphoramidon was added 30 minutes before the SIFa neuropeptide was added to the medium in which the cells were cultured. Cell supernatant was collected at 4 time points (0, 15 min, 30 min, and 60 min). Reverse-phase high-performance liquid chromatography (HPLC) was used to measure the cleavage of physiological peptides. **(D-F)** HPLC analysis revealed a progressive degradation of SIFa, which was significantly reduced in the presence of the Nep1 inhibitor. **(D)** Comparisons of mean progressive degradation analyzed after the indicated time (0, 15, 30, and 60 min). The bars represent the mean ± SE; n=3. **(E)** Plots represent mean ± SE, and each dot represents data measurement for one experiment. Unpaired t-tests were used between the two treatments. **(F)** Paired comparisons of progressive degradation were analyzed for each treatment between the indicated time (0 min) and the end of the assay at 60 minutes. Paired t-tests were used within the same treatment.

In a next step to test our hypothesis, we asked whether Nep1 encodes an active peptidase capable of cleaving SIFa. We first generated S2 cells stably expressing an inducible Nep1 construct (see Figure S2A for map of construct). Western blot analysis confirmed efficient induction of Nep1 expression, whereas uninduced cells and control S2 cells lacking the inducible construct showed no detectable Nep1 expression (see Figure S2B). We then added the selective Mca-RPPGFSAFK substrate at different time points following Nep1 induction and measured fluorescence emission as a readout for enzymatic activity. Nep1 induction resulted in robust enzymatic activity that was sensitive to thiorphan and phosphoramidon, two specific inhibitors of Nep1 (see Figure S2C). This demonstrated that the inducibly expressed Nep1 is biologically active. Moreover, given that phosphoramidon produced the strongest enzymatic inhibition, we used it in subsequent experiments.

To determine whether SIFa is a substrate of Nep1, supernatants were collected at different time points during the enzymatic assay and analyzed by HPLC (Figure 3C, for schematic representation). This demonstrated that SIFa is cleaved by the peptidase Nep1, as its levels decreased over time in presence of Nep1, whereas its degradation was attenuated in the presence of the Nep1 inhibitor (unpaired t tests: SIFa vs SIFa + inhibitor, 0 min *p* > 0.999; 15 min *p* = 0.206; 30 min *p* = 0.055; 60 min *p* = 0.019) (Figure 3D, E). Furthermore, comparisons of SIFa levels between two time points, 0 and 60 minutes, within the same (each?) treatment, revealed significant degradation in the absence phosphoramidon (paired t-test *p* = 0.033), whereas no significant degradation was detected in its presence (paired t-test *p* = 0.4501) (Figure 3D).

Together these results demonstrate that SIFamide is a substrate for the mushroom body expressed and enzymatically active peptidase Neprilysin 1.

### Mushroom body expressed Nep1 regulates food seeking behavior and feeding microstructure

Given the expression of biologically active Nep1 in the mushroom bodies and our demonstration that SIFa is a Nep1 substrate, we next hypothesized that Nep1 would impact food-seeking behavior and feeding microstructure. To determine the role of Nep1 in regulating feeding behavior, we first analyzed hunger-driven food-seeking behavior. Starved flies with a P-element insertion in the Nep1 gene, Nep1^Ey21255^ exhibited extended hunger-driven food-seeking behavior (Kruskal-Wallis: H = 84.26, df = 4, *p* < 0.001) (Figure 4A). To provide additional evidence that the behavioral alterations were caused by the P-element insertion, we generated and tested a Nep1 revertant line by precise transposon excision. The revertant flies displayed a feeding behavior indistinguishable from wild-type control flies, Canton-S or iso31 (Dunn’s post hoc: *p* = 0.23, *p* = 0.29, respectively) (Figure 4A). We further determined the impact of the P-element insertion and found that it leads to increased expression of Nep1 (Figure S3), an effect that is reversed in the P-element excision line. Previous studies have shown that flies rely on their sense of smell to locate the target (Gorter et al., 2016; Libert et al., 2007). Therefore, as a positive control, we tested mutant flies lacking Orco, the co-receptor for odorant receptors (ORs), which are known to be unable to detect odors. We found that Orco mutant flies were unable to locate yeast drops, exhibiting a phenotype indistinguishable from that of Nep1^Ey21255^ flies (Dunn’s post hoc: *p* > 0.99) (Figure 4A). In addition, we quantified the number of sips and activity bouts during feeding using the FlyPAD. Nep1^Ey21255^ flies exhibited significantly fewer sips (Figure 4B) and activity bout numbers (Figure 4C) compared to Nep1 revertant flies (sips: Mann-Whitney U test =2484, *p* = 0.0001; activity bout: Mann-Whitney U test =2461, *p* = 0.0001).

**Figure 4:**
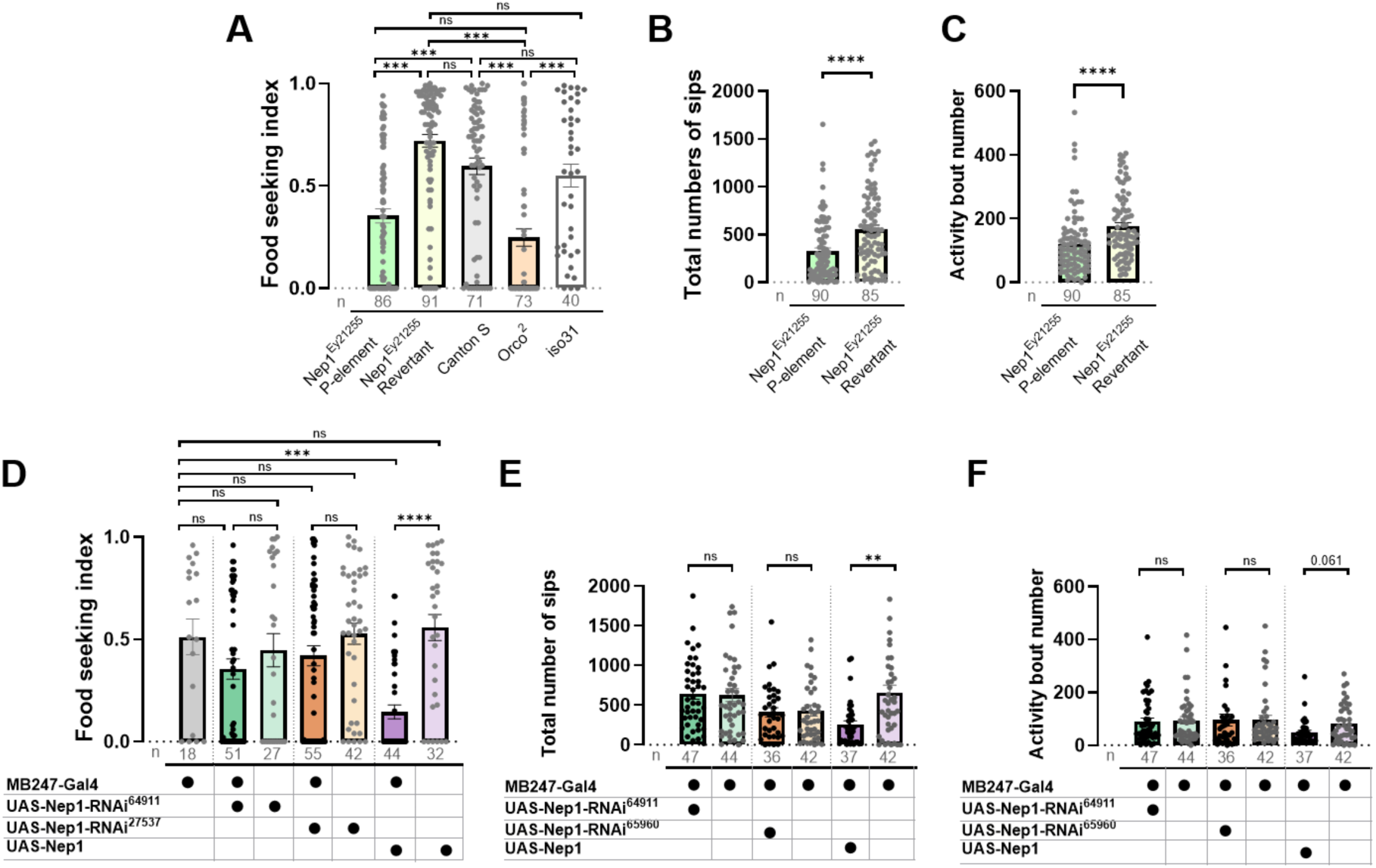
Overexpression of Nep1 in the mushroom body controls impaired-driven food-seeking behavior and food intake. **(A)** P-element insertion in Nep1EY21255 overexpression impairs food-seeking behavior. Hunger-driven food-seeking behavior in Nep1 revertant flies was indistinguishable from that of control flies. Orco mutant flies display food-seeking behavior like Nep1^Ey21255^ flies. Kruskal-Wallis tests followed by Dunn’s multiple comparisons test. **(B)** The total number of sips in Nep1 revertant flies was significantly higher than in Nep1^Ey21255^ flies. Mann-Whitney U test. **(C)** The activity boutnumberNep1 revertant flies was significantly higher than in Nep1^Ey21255^ flies. Mann-Whitney U test. **(D)** Knockdown of Nep1 in the MB did not significantly disrupt the feeding response. Overexpression of Nep1 in the MB decreased hunger-driven food-seeking behavior. Kruskal-Wallis tests followed by Dunn’s multiple comparisons test. **(E)** The total number of sips was not significantly different when Nep1 was knocked down in the MB. But overexpression of Nep1 in the MB reduced the number of sips compared to parental controls. Mann-Whitney U test. **(F)** The activity bout number was not significantly different when Nep1 was knocked down in the MB. Overexpression of Nep1 in the MB showed a strong trend toward reduced sips compared to parental controls. Mann-Whitney U test. Plots represent mean ± SE, and each dot represents a data measurement for one individual. The number of tested flies is represented below each bar for each genotype. Asterisks represent statistical significance: *p < 0.05, **p < 0.01, ***p < 0.001, ****p < 0.0001, ns, not significant.

The Nep1 P-element insertion line is expected to affect Nep1 expression in all cell types in which it is expressed. To determine whether the observed phenotypes could be recapitulated by selectively overexpressing Nep1 in the mushroom bodies, we used of the GAL4-UAS system and the mushroom body-specific GAL4 line MB247. Overexpression of Nep1 specifically in the MB significantly impaired hunger-driven food-seeking behavior (Kruskal-Wallis: H = 28.59, df = 2, *p* < 0.0001) (Figure 4D). In addition, Nep1 overexpression in the mushroom bodies resulted in a significant reduction in the number of sips and activity bouts (Figure 4E, 4F). Therefore, we can conclude that Nep1 in the mushroom bodies regulates both food seeking and feeding microstructure. In a reciprocal experiment, we examined the impact of Nep1 knockdown in the MB and found that it does not significantly affect food seeking behavior compared with the corresponding parental controls (Figure 4D) for either of the two Nep1-RNAi effector lines tested: 64911 (Kruskal-Wallis: H = 2.571, df = 2, *p* = 0.2766) and 27537 (Kruskal-Wallis: H = 2.793, df = 2, *p* = 0.2475). We also found that Nep1 knockdown in the MB did not significantly affect feeding microstructure compared with the corresponding parental controls for either of the Nep1-RNAi effector lines tested (Figure 4E and 4F).

## DISCUSSION

Hunger-driven food-seeking behavior is crucial for maintaining energy homeostasis and ensuring survival and encompasses localization and consumption of food resources when energy stores are low (Kelly et al., 2022; Lin et al., 2019; N. K. Smith & Grueter, 2022). These processes involve olfactory and gustatory detection of food cues, foraging and the microstructure of feeding. Hunger and satiety modulate these behaviors through various neural circuits and the integration of multiple signals that reflect feeding status. In *D. melanogaster*, numerous neuropeptides are involved in encoding hunger and satiety states and in regulating appetitive behavior, foraging and food intake, and metabolic homeostasis (Lin et al., 2019; Nagata & Zhou, 2019; Nässel, 2025; Nässel et al., 2019; Nässel & Zandawala, 2020; Yapici, 2025). An important question is how this information is integrated and how the activity of peptidergic neurons and of peptide hormones secreted into circulation is coordinated to elicit appropriate behavioral responses according to the internal state of the animal.

SIFamidergic neurons have emerged as central integrators of behavioral states (Zhang & Kim, 2026) with established roles in reproductive behavior (Sellami & Veenstra, 2015; Song et al., 2025; Terhzaz et al., 2007; Zhang et al., 2025), sleep (Cavanaugh et al., 2014; Huang et al., 2021; Park et al., 2014; Velazquez et al., 2025) and feeding behavior (Dreyer et al., 2019; Martelli et al., 2017; Velazquez et al., 2025; this study). These behaviors are mutually exclusive and thus require a dynamic and finely tuned regulation of SIFamidergic activity to balance competing behavioral states. This central integrator function is supported by the extensive projections of SIFamidergic neurons throughout the brain and by direct input from multiple neuronal populations (Cavanaugh et al., 2014; Dreyer et al., 2019; Martelli et al., 2017; Terhzaz et al., 2007). However, the downstream targets and mechanisms through which SIFamidergic neurons regulate behavior remain poorly understood. Known examples of downstream neurons include fruitless-expressing neurons in reproductive behavior (Sellami & Veenstra, 2015) and antennal lobe neurons involved in the detection of food-associated odors (Martelli et al., 2017). Further candidate downstream neurons were identified based on anti-SIFamide immunohistochemistry (Martelli et al., 2017) and Trans-tango based identification of synaptic contacts (Dreyer et al., 2019) and include neurons in the optic lobes, the suboesophageal ganglion, the dorsal fan-shaped body, and the mushroom bodies. However, the functional contribution of these synaptic partners downstream of SIFamidergic neurons has remained largely unexplored. Here, we show that SIFamidergic input to the mushroom bodies is direct. Although this input likely involves synaptic transmission, based on the results of the present study, we cannot exclude that volume transmission is also involved. The fact that selective downregulation of the SIFa receptor in the mushroom bodies strongly impaired food-seeking behavior and altered the microstructure of feeding, identify the mushroom bodies as a functionally relevant downstream target of SIFamidergic signaling. Additional indirect SIFamidergic may also contribute to these effects through the neural populations that we identified as being connected to the mushroom body circuitry. Consistent with a broader role of the mushroom bodies in integrating feeding-related signals, Tsao et al. (2018) previously demonstrated that four different neuropeptides (AstA, NPF, sNPF and DILPs) act on mushroom bodies through distinct types of dopaminergic neurons to regulate feeding behavior. Combined with the findings of the present study, this shows that the mushroom bodies integrate multiple feeding-relevant signals and contribute to the control of innate food-seeking behavior and foraging, in addition to their well-established role in olfactory conditioning and learning and memory (Akalal et al., 2006; Çoban et al., 2024; Heisenberg et al., 1985; Hige, 2018; Lin et al., 2014; McGuire et al., 2001; Turner et al., 2008).

Our findings further reveal that the mushroom bodies are not only downstream targets of SIFamidergic signaling but can also modulate SIFamide signaling through the peptidase Neprilysin 1. We show that SIFamide is a substrate for Nep1 and that increasing Nep1 expression in the mushroom bodies phenocopies the behavioral consequences of SIFaR loss, including impaired food seeking and disruption of feeding microstructure. These findings are consistent with increased degradation of SIFamide and a subsequent reduction in SIFamidergic signaling in vivo. Thus, we propose that Nep1 provides an additional layer of regulation of feeding behavior by modulating the local availability of SIFamide within the mushroom bodies. Interestingly, Hallier et al. (2016) have previously shown that Neprilysin 4 regulates food intake and insulin-like peptide expression. They showed that this effect is mediated by peptide substrates of Neprilysin 4, with established roles in feeding and the regulation of ILP expression (Hallier et al., 2016). These authors also observed that, contrary to overexpression, knockdown of Nep4 had only very limited effects. We also saw only mild effects of knockdown of Neprilysin 1. One possible explanation for this is functional compensation by other neprilysins. This possibility was suggested by Hallier et al. (2016) for Neprilysin 4. However, compensation appears less likely in the case of Nep1, given that, to our knowledge, Nep1 is the only neprilysin expressed in the mushroom bodies.

Martelli et al. (2017) previously showed that SIFamidergic neurons promote appetitive and gustatory behavior. Our findings that the mushroom bodies constitute a direct downstream target of SIFamidergic signaling to regulate food seeking and feeding microstructure provides further support for the notion that SIFamidergic neurons act as a central integrator as suggested by Zhang and Kim (2026). Given that SIFamide signaling also regulates sleep, it will be interesting to determine whether targeted manipulation of the SIFa receptor in the dorsal fan-shaped body, a neuropil previously shown to be important for sleep (Donlea et al., 2011) will disrupt that behavior. More broadly, our findings support a model in which SIFamidergic neurons directly innervate all relevant neuronal populations for the mutually exclusive behaviors sexual reproduction, sleep and hunger. Aggression represents another behavior that is mutually exclusive with sexual reproduction and sleep and can itself be induced by starvation (Anholt & Mackay, 2012; Zwarts et al., 2011). Given that injection of SIFamide leads to increased aggressive behavior in the freshwater prawn *Macrobrachium rosenbergii* (Vázquez-Acevedo et al., 2009), it will be interesting to determine whether SIFamide also contributes to the regulation of aggression in *Drosophila*.

Finally, SIFamide is conserved across arthropods suggesting an evolutionarily ancient association with feeding-related behaviors (Arendt et al., 2016; Ayub et al., 2020; Christie et al., 2006; Janssen et al., 1996; Martelli et al., 2017; Ons et al., 2011; Terhzaz et al., 2007; Verleyen et al., 2004, 2009). Mushroom body-expressed genes have been shown to be important for shaping the evolution of foraging behavior in Drosophila species (Chen et al., 2012). We propose that some of these genes may influence foraging behavior by modulating SIFamidergic signaling. In this context, the observation that Neprilysin 1 is expressed in mushroom body neurons in nine drosophilid species of the melanogaster species subgroup (Rebeiz et al., 2011), raised the possibility that the SIFa-Nep1 relationship is also evolutionarily conserved. This hypothesis remains to be tested. More broadly, the recruitment of additional genes into the regulation of SIFamidergic signaling could provide an additional layer of regulation through which species-specific behavioral responses are shaped, particularly if the expression or activity of these genes is itself regulated by internal state. Overall, our findings suggest the evolutionarily ancient SIFamidergic signaling system can be fine-tuned by a peptidase and provide a potential mechanism through which the recruitment of novel regulatory genes may contribute to the diversification of feeding behavior.

## ACKNOWLEDGMENTS

Research at BF2i was supported by the Institut National de la Recherche pour l’Agriculture, l’Alimentation et l’Environnement (INRAE) and the Institut National des Sciences Appliquées de Lyon (INSA Lyon). P.C. was supported by FWO grants G065408.N10, G078914N, and by KULeuven grants C14/17/099 and C14/22/126.

## AUTHOR CONTRIBUTIONS

Conceptualization: E-S.B, P.C; Investigation: E-S.B., D.M., M.P.V., V.V., S.D.G., I.R., J.Z.; Analysis: E-S.B., D.M., M.P.V., V.V., S.D.G., I.R., J.Z., P.D.S., T.R., F.C., P.C.; Resources: E-S.B., D.M., V.V., P.D.S., T.R., F.C., P.C.; Supervision: E-S.B., V.V., P.D.S., T.R., F.C., P.C.; Funding acquisition: D.M., P.D.S., T.R., F.C., P.C.; Writing – original draft: E-S.B., D.M., M.P.V., T.R., P.C.; Writing – review and editing: E-S.B., D.M., M.P.V., V.V., S.D.G., I.R., J.Z., P.D.S., T.R., F.C., P.C.

