## Supplemental Figures for "Neprilysin 1 in the Mushroom Body controls Feeding behavior by regulating SIFamide levels in *Drosophila melanogaster*"

El-Sayed Baz<sup>1,2,#</sup>, Daan Moons<sup>1,#</sup>, Maria Pons Vizcarra <sup>1,#</sup>, Veerle Vulsteke<sup>1</sup>, Sofie De Groef<sup>1</sup>, Isabelle
6 Rahioui<sup>3</sup>, Jiajun Zhang<sup>4</sup>, Pedro Da Silva<sup>3</sup>, Thomas Riemensperger<sup>4</sup>, Federica Calevro<sup>3</sup>, Patrick
7 Callaerts<sup>1\*</sup>
8

Supplementary figures

Figure S1.

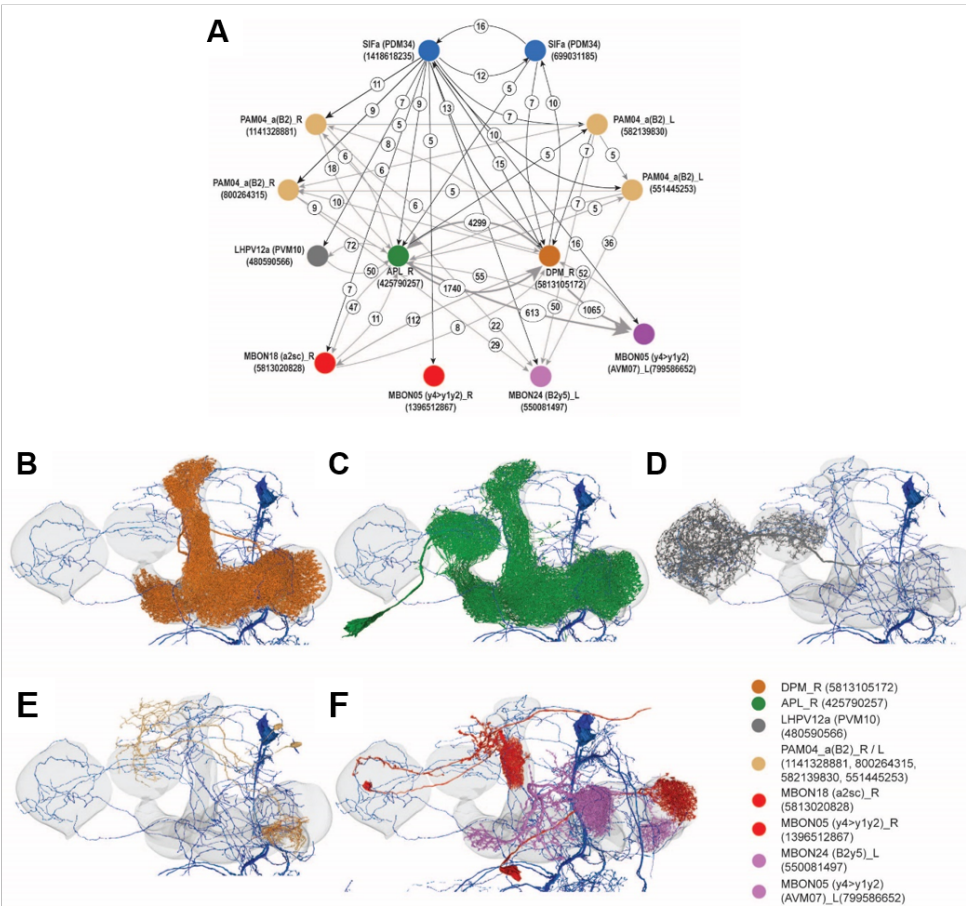

Figure S1: Interaction of SIFamidergic neurons and mushroom body circuits:

(A) Connectome of SIFa - Mushroom body circuits: synaptic connectivity of SIFamidergic neurons to mushroom body circuit neurons (synaptic output is indicated by the direction of the arrow, synaptic weight by the numbers in circles, direct connectivity between SIFamidergic neurons and MB circuit neurons is indicated by black arrows, connectivity among MB circuit neurons not interacting with SIFamidergic neurons are indicated in grey). SIFa neurons show synaptic interaction among each other and to APL (C, green), DPM (B, orange), PAM04 dopaminergic neurons (E, yellow) innervating beta 2 lobes of the mushroom, lateral horn output neurons (D, grey) and mushroom body output neurons from the compartments $\gamma 1, 2$, $\gamma 4$, $\gamma 5$, $\alpha 2$, (F, red, magenta).

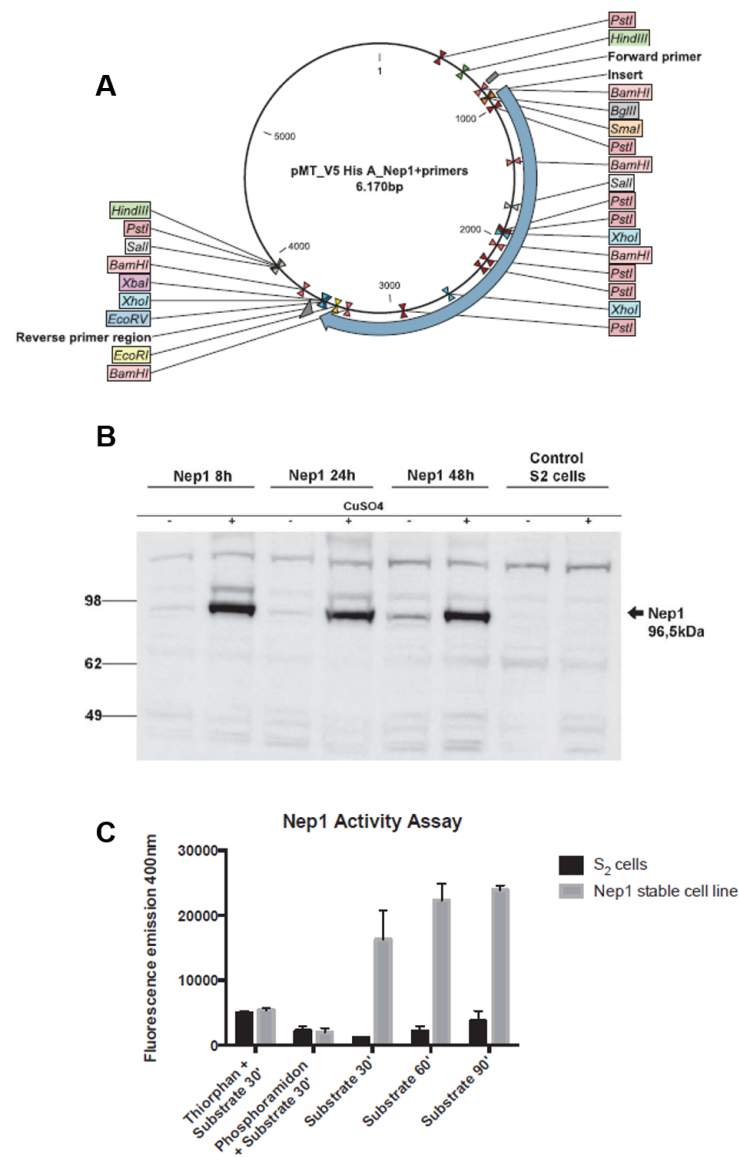

**FigureS2: Expression of Nep1 in *Drosophila* Schneider 2 (S2) cells.**

**(A)** Vector pMT\_V5 His A\_Nep1 transfected into S2 cells. **(B)** Western Blot showing Nep1 expression in *Drosophila* S2 cells. Cells were lysed after the respective induction times; S2 control cells were harvested after 24 hours. 25 µg of protein was loaded in the gel. Rabbit anti-Nep1 (E1) (IgG mix) was used as primary antibody and goat anti-rabbit HRP was used as secondary antibody. Nep1 protein is expected to be around 96,5 kDa. **(C)** Neprilysin 1 activity assay. Around 6 million S2 cells and Nep1 stable cell line were used for each assay. The inhibitors used were thiorphan and phosphoramidon, which were incubated 30min before adding the substrate. The substrate was added, the supernatant was collected after the indicated time (30, 60 or 90 min) and the fluorescence emission was analyzed. The bars represent the mean of the emission ± SEM; n=3

**Figure S3**

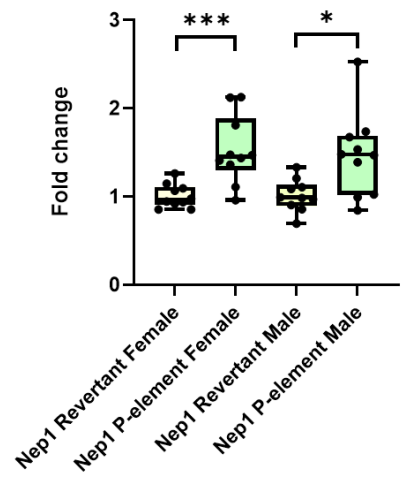

36
**Figure S3: qRT-PCR analysis of Nep1 expression in the Nep1<sup>EY21255</sup> and revertant lines.**
